# No evidence from complementary data sources of a direct projection from the mouse anterior cingulate cortex to the hippocampal formation

**DOI:** 10.1101/2022.01.25.477805

**Authors:** Lilya Andrianova, Steliana Yanakieva, Gabriella Margetts-Smith, Shivali Kohli, Erica S Brady, John P Aggleton, Michael T Craig

## Abstract

The connectivity and interplay between the prefrontal cortex and hippocampus underpin a number of key cognitive processes, with changes in these interactions being implicated in both neurodevelopmental as well as neurodegenerative conditions. Understanding the precise cellular connections through which this circuit is organised is, therefore, vital for understanding these same processes. Overturning earlier findings, a recent study described a novel excitatory projection from anterior cingulate cortex to hippocampus. We sought to validate this unexpected finding using multiple, complementary methods: anterograde and retrograde anatomical tracing, using both anterograde and retrograde AAVs and monosynaptic rabies tracing. Additionally, an extensive data search of the Allen Projection Brain Atlas database was conducted to find the stated projection within any of the deposited anatomical studies, as an independent verification of our own results. However, we failed to find any evidence of a direct, monosynaptic projection from mouse anterior cingulate cortex to the hippocampus proper.

## Introduction

Models of the functional interactions between the rodent prefrontal cortex and the hippocampal formation have had to address their asymmetrical relationship. The subiculum and CA1 give rise to direct efferents that terminate in the prelimbic, infralimbic, and medial orbital cortices (Cenquizca and Swanson, 2007; Groenewegen et al., 1987; Jay and Witter, 1991; Swanson, 1981) as well as light projections to the anterior cingulate cortex (Cenquizca and Swanson, 2007; Jay and Witter, 1991). In contrast, direct projections from the prelimbic and anterior cingulate cortices to the CA fields or the subiculum are not observed (Jones and Witter, 2007; Segal and Landis, 1974; Sesack et al., 1989; Vogt and Miller, 1983). Meanwhile, only a very sparse projection from infralimbic cortex to CA1 has been described (Hurley et al., 1991), with a recent preprint providing evidence of long-range inhibitory projections from prelimbic and infralimbic cortices to hippocampus proper (Malik et al., 2021). For this reason, models of prefrontal regulatory action upon the hippocampus have emphasised the importance of indirect routes. These routes include relays via parahippocampal cortices and via subcortical sites such as nucleus reuniens and the anterior thalamic nuclei (Eichenbaum, 2017; Furtak et al., 2007; Jones and Witter, 2007; Prasad and Chudasama, 2013).

This relationship was transformed by the description of light, direct projections from the dorsal anterior cingulate cortex to the CA1 and CA3 fields in mice (Rajasethupath et al., 2015). Despite their sparsity, optogenetic manipulation of these same projections was sufficient to elicit contextual memory retrieval, and optogenetic stimulation of these axons evoked robust excitatory post-synaptic currents in pyramidal cells in both CA3 and CA1 (Rajasethupath et al., 2015). These findings are striking, not least because by revealing a previously unknown connection in the rodent brain, they suggest that there are potentially many such functional connections still waiting to be uncovered.

Given the impact of their functional findings on anterior cingulate efferents (Rajasethupath et al., 2015), the present study re-examined the status of the projections from the anterior cingulate area (ACA) to the hippocampal formation (dentate gyrus, CA fields, and subiculum) in mice. Anterograde tracing using adeno-associated viral vectors (AAVs) found no evidence of the anterior cingulate to hippocampus projection in mice, nor did rabies virus-assisted monosynaptic retrograde tracing from hippocampal pyramidal cells. Furthermore, a thorough data mining analysis of Allen Expression data also revealed no evidence of an anterior cingulate to HPC projection in mice.

## Methods

All UK-based research was carried out in accordance with the UK Animals (Scientific Procedures) Act 1986 and was subject to local ethical review by the Animal Welfare and Ethical Review Board at the University of Exeter. All animals were maintained on a 12 h constant light / dark cycle and had access to food and water *ad libitum.* We used standard enrichment that included cardboard tubes, wooden chew blocks and nesting material.

### AAV anterograde and retrograde tracing

This tracing study used adult C57BL/6 or *SST*-Cre mice of both sexes aged 2 to 5 months, ranging from 22.2 g to 31.3 g (mean age 3 months, mean weight 25 g). To target the ACA, we made stereotaxic injections of 250 nl of AAV5-CamKII-GFP 5.3×10^12^ vg/ml (Viral Vector Facility, Neuroscience Centre Zurich, Swizterland) or AAV5/2-hSyn1-ChR2_mCherry 7.1×10^12^ vg/ml (Viral Vector Facility, Neuroscience Centre Zurich, Switzerland). Briefly, mice were anaesthetised with 5% isoflurane, were placed on a heated pad for the duration of the surgery, and maintained at 1.5 to 2.5% isoflurane (with a flow rate of ~2 Lmin^-1^ O2). Mice were given 0.1 mg/kg of buprenorphine (buprenorphine hydrochloride, Henry Schein) subcutaneously at the start of surgery as an adjunct analgesic, and carprofen (Rimadyl, Henry Schein) was given at a dose of 5 mg/kg subcutaneously at the end of surgery and on subsequent days, as required. An incision was made down the midline and a craniotomy was performed to allow injection of virus (250 nl). For anterograde tracing, we first used the co-ordinates reported by Rajasethupathy et al., 2015: A/P +1.0 mm (relative to Bregma), M/L −0.35 mm and D/V-1.2 mm (from pia). Additionally, to minimise spread of virus into M2 cortical area, we also injected at A/P +1 mm (relative to Bregma), M/L −0.2 mm and D/V-1.3 mm (from pia) on the same rostrocaudal plane, and targeted ACA at a more rostral site A/P +1.75 mm (relative to Bregma), M/L −0.2 mm and D/V-1.5 mm (from pia). For retrograde tracing with AAVs, we used rAAV-retro, developed at HHMI Janelia Campus (Tervo et al., 2016), and injected rAAV-retro/2-CAG-EGFP_Cre-WPRE-SV40p(A) 5.9 x 10^12^ vg/ml (Viral Vector Facility, Neuroscience Centre Zurich, Switzerland) into hippocampal region CA1, using the following co-ordinates: A/P −2 mm (relative to Bregma), M/L −1.5 mm and D/V-1.35 mm (from pia) After surgical repair of the wound, mice were given 5 mg/kg carprofen (Rimadyl, Henry Schein) subcutaneously immediately after surgery, and again the following day. Further analgesia was provided as required. The mice were maintained for at least 3 weeks to provide sufficient time to expression of virally-delivered transgenes, and were killed by transcardial perfusion / fixation with 4% paraformaldehyde (Sigma-Aldrich, UK) in 0.1 M phosphate buffer.

Perfused brains were cryoprotected with 30% sucrose solution (Fisher Schientific, UK) and sliced at 30 μm using a SM2010R freezing microtome (Leica, UK). The slices were stained with DAPI (HelloBio, UK) and fluorescence was visualised using CoolLED pE-4000 (CoolLED, UK); photos taken on Nikon 800 microscope. For anterograde tracing experiments 32 mice were injected (2 were excluded due to failed injections as no fluorescence was observed) and 10 mice were injected with a retrograde virus (2 mice were excluded due to failed injections). Viral expression patterns in individual mice at the injection sites of both sets of coordinates were overlayed (figure 1A-B) to show the variation in the injections and typical spread of the virus in the injection areas. A qualitative assessment of the projection patterns was performed and representative images of regions of interest are presented on figure 1C-I.

**Figure 1:**
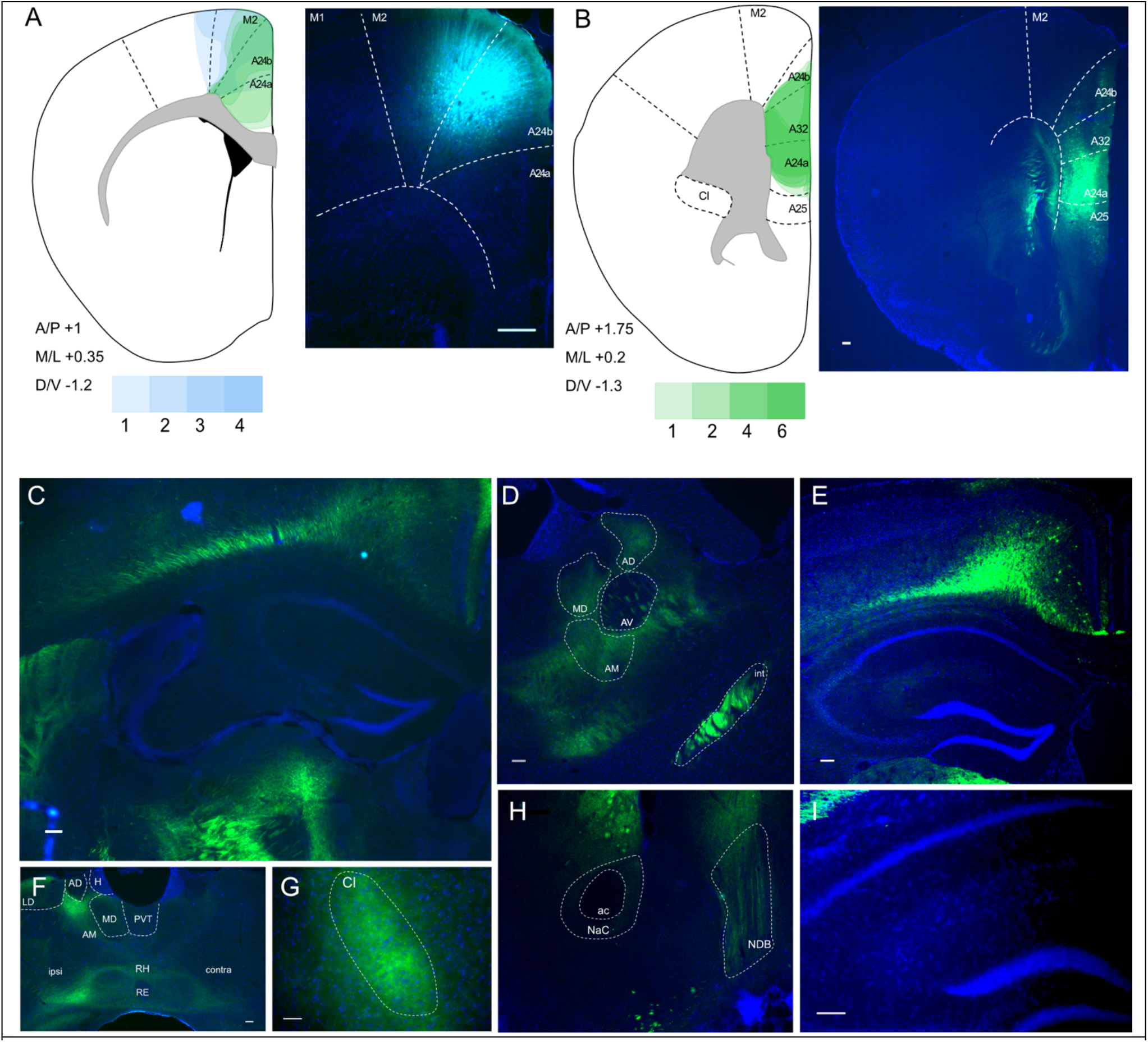
Anterograde viral tracing from ACA CaMKII-expressing neurons shows no excitatory projection to the dCA1 region of hippocampus. **(A)** Prefrontal cortex injection sites and spread, blue overlays show injections made using previously published coordinates (Rajasethupathy et al., 2015), green overlays show in-house initial optimization attempt, n= 3 and n=3 respectively, **(B)** Prefrontal cortex injection site and spread of individual injections using in-house optimized coordinates; each overlay shape represents one injection, n=7. **(C)** Representative images of the regions of interest and known target regions of the PFC, **(D)** ipsilateral thalamic nuclei and internal capsule, **(E)** dorsal hippocampus shows no fibres, unlike the overlaying RSC and other cortices, **(F)** rostral and midline thalamus in both ipsi and contralateral hemispheres, showing both fibre and terminal label, **(G)** fibres in ipsilateral claustrum, **(H)** fibres are found in diagonal band nucleus and striatum, but absent in nucleus accumbens, **(I)** higher magnification image of dorsal CA1 region of the hippocampus, showing no fibres. A24a – area 24a (infralimbic), A24b – area 24b (cingulate cortex 1), A25 – area 25 (dorsal peduncular cortex), A32 – area 32 (prelimbic), ac – anterior commissure, ACA – anterior cingulate area, AD – anterodorsal nucleus, AM – anteromedial nucleus, AV – anteroventral nucleus, Cl – claustrum, H – habenula, int – internal capsule, LD – lateral dorsal nucleus, M1 – primary motor area, M2 – secondary motor area, MD – mediodorsal nucleus, NaC – nucleus accumbens core area, NDB – diagonal band nucleus, PVT – paraventricular nucleus of the thalamus, RE – nucleus reuniens, RH – rhomboid nucleus. Scale bar 100 microns

### Monosynaptic Retrograde tracing

For monosynaptic rabies tracing, we used methods reported by others (Sun et al., 2014). Adult *Emx1-cre* mice (Guo et al., 2000) were crossed with floxed TVA mice (Seidler et al., 2008) to allow specific targeting of pyramidal cells; *Emx1-cre* mice alone were used as controls to ensure the rabies virues did not transduce neurons in the absence of TVA gene. Mice of both sexes aged 3 to 7 months, ranging from 21.7 g to 40.6 g (mean age 4.5 months, mean weight 27.2 g). To target the efferent projections to dCA1, we made stereotaxic injections of AAV8-FLEX-H2B-GFP-2A-oG, titre 3.93×10^12^ vg/ml (Salk Institute Viral Vector Core) followed by injection EnvA G-deleted Rabies-mCherry 6.13×10^8^ vg/ml (Salk Institute Viral Vector Core or Charité – Universitätsmedizin Berlin Viral Vector Core) 2 weeks after the initial viral injection. Briefly, mice were anaesthetised with 5% isoflurane, were placed on a heated pad for the duration of the surgery, and maintained at 1.5 to 2.5% isoflurane (with a flow rate of ~2 Lmin^-1^ O2). Mice were given 0.1 mg/kg of buprenorphine (buprenorphine hydrochloride, Henry Schein) subcutaneously at the start of surgery as an adjunct analgesic, and carprofen (Rimadyl, Henry Schein) was given at a dose of 5 mg/kg subcutaneously at the end of surgery and on subsequent days, as required. An incision was made down the midline and a craniotomy performed to allow injection of virus (250 nl). We targeted hippocampal region CA1 at dorsal and ventral points, using the following co-ordinates: A/P −2 mm (relative to Bregma), M/L −1.5 mm and D/V-1.35 mm (from pia) and A/P −2.8 mm (relative to Bregma), M/L −2.4 mm and D/V-4.2 mm (from pia) for dorsal and ventral CA1 regions respectively. After surgical repair of the wound, mice were given 5 mg/kg carprofen (Rimadyl, Henry Schein) subcutaneously immediately after surgery, and again the following day. Further analgesia was provided as required. The mice were maintained for 2 weeks to provide optimal time for expression, and were killed by transcardial perfusion / fixation with 4% paraformaldehyde (Cat number P6148 Sigma-Aldrich, UK) in 0.1 M phosphate buffer.

Perfused brains were cryoprotected with 30% sucrose solution (Fisher Schientific, UK) and sliced at 50 μm using a SM2010R freezing microtome (Leica, UK). The slices were mounted using HardSet Mounting Medium with DAPI (Vector Labs) and fluorescence was visualised using CoolLED; photos taken on Nikon 800 microscope. A total of 18 mice including 4 controls were injected (2 were excluded due to failed injections as no fluorescence from both viral constructs was observed). Representative images of the spread of the viruses can be found in figure 2. A semi-quantitative assessment of the projection patterns was performed by recording the number of slices with the cells expressing the virus in each region of interest present and the summary of this data is shown (figure 2P). These monosynaptic rabies tracing data were generated in the same experiment that we have reported elsewhere, focusing on connections between nucleus reuniens and CA1 pyramidal cells (Andrianova et al., 2021).

**Figure 2:**
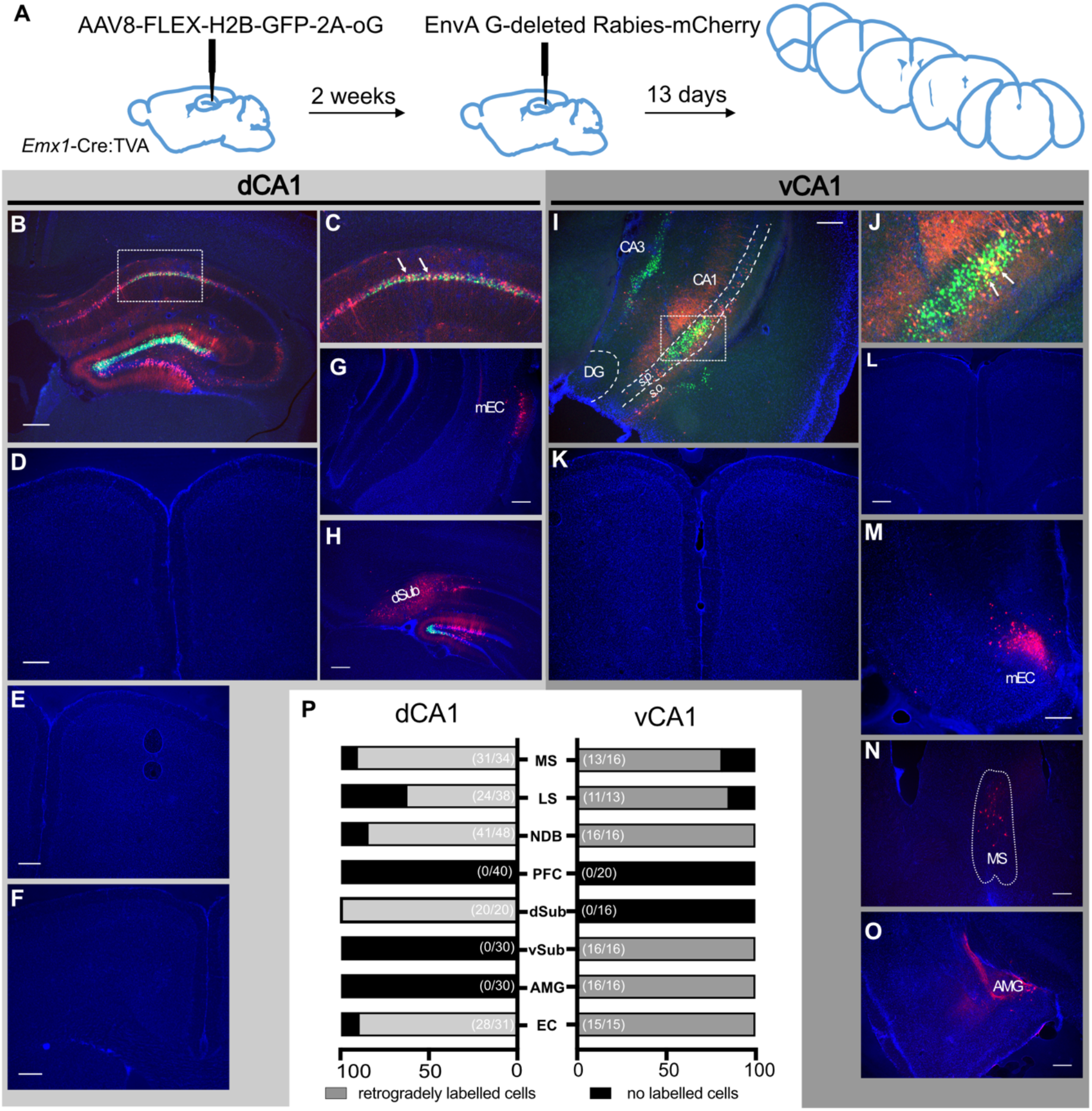
Monosynaptic retrograde tracing from dCA1 and vCA1 using modified rabies virus shows no excitatory projection from the ACA to either dorsal or ventral CA1 regions. **(A)** outline of experimental method. **(B,C)** representative images of dCA1 injection site, white arrows show starter cells. Representative images of: **(D-F)** prefrontal cortical areas at +1.8, +1.5 and +1.1 Bregma, **(G)** entorhinal cortex, **(H)** dorsal subiculum. **(I, J)** vCA1 injection site, white arrows show starter cells. Representative images of: **(K,L)** prefrontal cortical areas at +1.5 and +1.2 Bregma, **(M)** entorhinal cortex, **(N)** medial septum, **(O)** amygdala. **(P)** Summary of data. Bar graphs show the averge percentage of sections from each animal showing presynaptic neurons while numbers in bracket show total number of sections examined. Scale bar represents 100 micron.

### Allen Projection Atlas Data Mining

The *Allen Mouse Brain Connectivity Atlas* (http://mouse.brain-map.org/) is a comprehensive database of images of axonal projections in the mouse brain. To create the atlas, each mouse brain is injected with enhanced green fluorescent protein (EGFP) expressing adeno-associated virus (AVV) as an anterograde tracer into a *source* brain region (for further details see http://connectivty.brain-mpa.org). Then, the axonal projections are systematically imaged using a TisseCyte 1000 serial two-photon tomography system. The data in the atlas were collected from adult mice on postnatal day P56 ± 2 (for details, see Oh et al., 2014). Each case in the atlas contains high resolution images and quantified projection information based on the optical density of label. Detailed histograms of the signal in each structure are presented giving the projection volume (mm^3^) and projection density (the fraction of area occupied by a fluorescent signal from the viral construct used relative to the whole structure). Details of the algorithms used are provided (Kuan et al., 2015). The sensitivity of this viral tracer has been compared with biotinylated dextran amine (BDA), revealing comparable transport properties (http://connectivty.brain-mpa.org).

To establish whether there are direct projections from the anterior cingulate area (dorsal ACA and ventral ACA) to the hippocampal formation [Cornu Ammonis (CA) 1-3, dentate gyrus (DG), and subiculum (SUB)], a comprehensive systematic search was completed. In addition, the density of fibre label in the cingulum bundle (CB) was taken from the Allen Atlas. The dorsal and ventral ACA were entered as a *source structure* and the CA1-3, DG, SUB, and cingulum bundle as target structures. The search was further filtered for: (1) Reporter type: EGFP; (2) Hemisphere: Either; (3) Minimum target injection volume: 0.0001 mm^3^. A total of 99 cases were returned. Then, 63 cases with injection volumes of 0.02 – 0.3mm^3^ in the ACA were selected for further processing, of which 47 had a minimum of 90% of the injected virus (mean = 0.108 mm^3^; *SD* = 0.059 mm^3^) within the dorsal and ventral anterior cingulate cortices. Table 1 lists all 47 cases and provides further information about the transgenic lines and the individual injections. The large majority (n = 44) of the injections were made in the right hemisphere, the remaining three in the left. In the few cases where some signal was reported in areas CA1 and CA3, we used the Willcoxon signed rank test to compare that signal with the corresponding signal in the ipsilateral DG and CA2 to help test for background variation. (The dentate gyrus and CA2 were selected as there are no reports of ACA inputs to these subareas.)

There were three cases of wild-type mice (C57BL/6J) with 100% of the injection volume within ACA. Of the 44 transgenic mice, 28 had 100% of the tracer within the ACA, making a total of 31 such cases. The other 16 cases had a minimum of 90% of the tracer in the ACA and the remainder in adjacent areas, including the secondary motor area, the prelimbic cortex, and adjacent fibre tracts in various proportions. These 16 cases were, therefore, not considered in the initial analyses. In the WT group all mice were male, whilst in the transgenic group of 28 cases with 100% of the injection in the ACA, 12 were female. In the same transgenic group, 27 of the 28 injections were in the right hemisphere (see Table 1 for a list of included studies).

## Results

### Anterograde virus-assisted tracing

We carried out stereotaxic injections of AAV vectors to allow expression of GFP or mCherry in ACA neurons to allow us to map their projections (figure 1). We targeted our injections into ACA / mPFC at two different AP levels: 1.0 mm rostral to Bregma (figure 1A) and 1.75 mm rostral to Bregma (figure 1B). The more caudal co-ordinates exactly matched those reported by Rajasethupath and colleagues (2015), which also produced some somatic labelling in M2. We then targeted a site deeper and slightly more lateral within a more rostral point in the rostrocaudal axis, which led to somatic labelling restricted entirely within the subdivisions of the prefrontal cortex. In all cases, we saw prefrontal efferents within retrosplenial cortex and several thalamic nuclei, as expected, but found no evidence of axons in any hippocampal subfield or subiculum (figure 1C, E, I). A summary of the projections from our anterograde labelling experiment is presented in supplementary Table 1, which incorporates data from a total of 30 mice. Supplementary figure 1 presents more examples from the anterograde tracing experiments.

### Monosynaptic retrograde tracing

After examining with anterograde virus-assisted tracing, we next used mononsynaptic rabies-assisted viral tracing to determine whether pyramidal cells in hippocampal region CA1 received direct inputs from ACA (experimental protocol summarised in figure 2A). We carried out tracing from both dorsal CA1 (dCA1; n = 6 mice; figure 2B – C) and ventral CA1 (vCA1; n = 4 mice; figure 2I - J). Acting as positive controls, and as we reported previously (Andrianova et al., 2021), retrogradely-labelled neurons were detected in all brain regions in all mice that one would expect to project to CA1: both dCA1 and vCA1 received monosynaptic inputs from medial vCA1 (figure 2M). As reported by others (Sun et al., 2014), we found evidence of monosynaptic inputs to dCA1 from dorsal subiculum (figure 2F) and to vCA1 from ventral subiculum, although we did not determine whether these neurons were GABAergic or glutamatergic. As expected, vCA1, but not dCA1, received monosynaptic input from the amygdala (figure 2L). Importantly, however, we found no evidence of monosynaptic projections from ACA or any other prefronal region to either dCA1 (figure 2D, G – H) or vCA1 (figure 2K, N – O). These data are summarised in figure 2P.

### Allen Atlas Data

A total of 31 cases had viral injections confined within the ACA. For these cases we recorded the optical density measures for transported fluorescent fibres in the regions of interest. The medians *(Mdn)* and ranges for each hippocampal region by group (WT or transgenic) are presented in figure 3. In the WT group (figure 3A and D) there was no evidence of signal in either the ipsilateral or contralateral CA1 field (all optical density measures = zero), and only one case contained any signal in the ipsilateral CA3 field (0.0002). For the transgenic group (figure 3B – D), the median signal for CA3 and CA1 remained at zero, but there were eight cases with potential signal in the ipsilateral and/or the contralateral CA1 and/or CA3 fields (all <0.0007). However, in all of these cases there was a low, but variable, background signal. For this reason, we compared the signal in areas CA1 and CA3 (which have potential anterior cingulate inputs) with that in the DG and area CA2 (neither of which are thought to receive anterior cingulate inputs). To maximise the likelihood of finding evidence for a projection we just used the eight cases with possible CA3 or possible CA1 label for the statistical comparisons.

**Figure 3:**
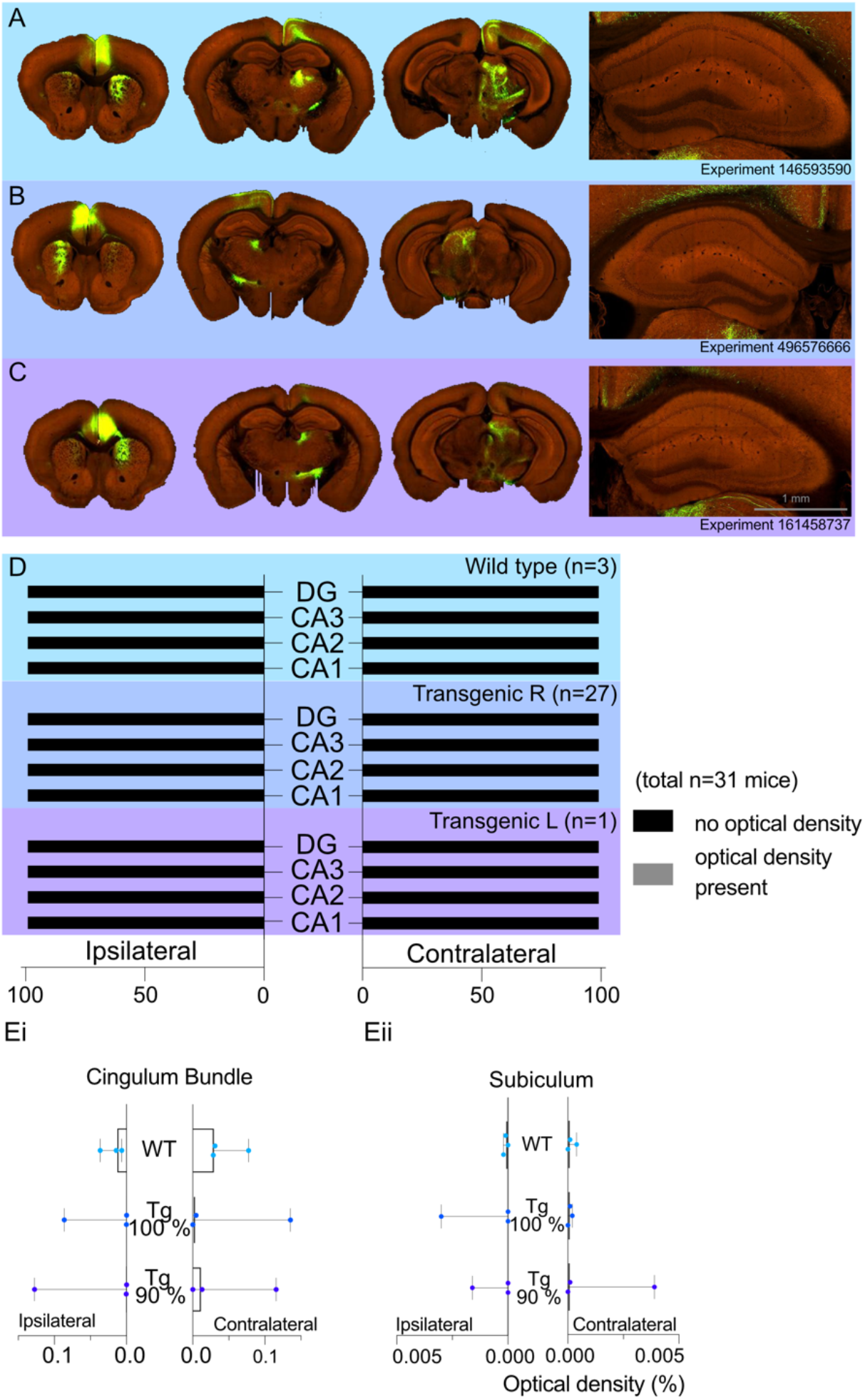
Allen Projection Atlas data search shows no evidence of a direct projection from anterior cingulate area to dorsal CA1. Representative examples of tracing experiments in **(A)** wild-type mouse line (C57BL/6J), **(B)** transgenic (Rbp4-Cre_KL100) mouse line experiment with 100% of the virus injected in the ACA, **(C)** transgenic (Rbp4-Cre_KL100) mouse line experiment with 90% of the virus injected in the ACA; and the corresponding typical anterograde expression patterns. **(D)** Histograms showing the summary of fluorescent fibres present in different hippocampal subfields in all datasets. No fibres were found any sections containing hippocampal regions DG and CA1-3. (**E**) Median signal densities of fibres seen in cingulum bundle (Ei) and subiculum (Eii) in experiments with wild type and transgenic mouse lines. The histograms show the optical densities of potential transported tracer in different experiments in the cingulum bundle and subiculum. While there is no apparent signal in the subiculum, the tracer uptake is significantly higher in the cingulum bundle. Of concern was whether the adjacent cortical field M2 might project to the hippocampal formation and, thereby, be a potential confound for tracer injections that extended into this area. For this reason we examined the *Allen Mouse Brain Connectivity Atlas* (http://mouse.brain-map.org/) for wild type cases meeting the inclusion criteria with injections of 90% or more within area M2. Any case in which the tracer extended into the anterior cingulate cortices was excluded. The search returned only one such case (ID: 585025284), which showed no evidence of a projection to CA1, CA2, or subiculum (all signal density counts zero). There was a very low signal count within the ipsilateral CA3 and DG (<0.0002). In an additional case (ID:141602484) the viral injection involved both the primary motor cortex (46%) and area M2 (54%). The only hippocampal signal was in the ipsilateral CA3 (<0.0002). Both cases had appreciably more signal in the ipsilateral than contralateral cingulum bundle, consistent with viable transport.

There was no evidence that either CA1 or CA3 contained higher signal levels in these eight cases, i.e., the higher optical measures reflected a higher background signal. Wilcoxon signed rank tests found no statistical difference in the signal between the ipsilateral DG and CA3 (*z*=-0.845, *p* = 0.388), CA2 and CA3 (*z*=-0.510*, p*=0.610), or the contralateral DG and CA3 (*z*=-0.755, *p*=0.438), and contralateral CA2 and CA3 (*z*=-1.841, *p* =0.066). Likewise, there were no significant differences between the signal in ipsilateral CA2 and CA1 (*z*=-1.63, *p*=0.102) or the contralateral CA2 and CA1 (*z*=-1.07, *p*=0.285). While there were significant differences between both the ipsilateral DG and CA1 (*z*=-2.4, *p*=0.016) and the contralateral DG and CA1 (*z*=-2.2, *p*=0.028), inspection of the data showed that in seven of the cases the signal was higher in the DG than CA1, i.e., the opposite direction to that consistent with an ACA projection to CA1.

In contrast, the signal in the cingulum bundle (both ipsilateral and contralateral) was higher in 29 of the 31 cases than the signal in CA1-CA3 and the DG. In the WT group the median signal density in the ipsilateral CB was 0.031 and 0.0014 in the contralateral CB 0.014 (see figure 3Ei for ranges). Similarly, the transgenic group with right-hemisphere injections had a median signal density in the ipsilateral CB of 0.0043 and 0.0002 in the contralateral CB (figure 3Ei). The presence of CB fibre label helps to confirm the effective transport of the tracer from the injection site. Lastly, there was no evidence of a projection from the ACA to the subiculum in any group as the median signal density observed for the ipsilateral and contralateral subiculum was a maximum of 0.0001 (Figure 3Eii).

The median density signal in the 16 transgenic cases with injection volumes of over 90% within the ACA was consistent with the data from the other 28 transgenic cases (see supplementary Table 1). The median group signal in both CA1 and CA3 was zero, with 10 cases having potential evidence of a signal (*max* = 0.0007) in CA1 and/or CA3. Again, Wilcoxon signed rank tests revealed no evidence of a statistical difference in the signal between the ipsilateral or contralateral DG and CA3 or CA2 and CA3 (*all z* <-1.63, *all p*s>0.102) nor between the ipsilateral and contralateral CA2 and CA1 (*all z* <-1.34, *all p_s_* >0.180). There were, however, statistically significant differences between both the ipsilateral DG and CA1 (*z*=-2.56, *p*=.010) and the contralateral DG and CA1 (*z*=-2.03, *p*=.042), but again the higher signal was in the DG. Further inspection of individual cases showed that one case (ID:125801033) had noticeably higher signal density in the ipsilateral DG (0.0108) and SUB (0.0039). A second case (ID: 585911240) had the highest signal density in the ipsilateral CA3 (0.0007). However, in both cases 5% of the injection had leaked into adjacent fibre tracts.

### Retrograde tracing using a single virus strategy

Thus far, we have failed to detect evidence of a direct projection from the cingulate cortex to hippocampus. The question remains of why there is a discrepancy between our data and that reported previously (Rajasethupathy et al., 2015). In our rabies-virus assisted retrograde tracing experiment, we used a pseudotyped glycoprotein-deleted rabies virus that could only infect neurons expressing TVA, and could only spread transynaptically from neurons that also expressed the rabies glycoprotein, delivered via a secondary helper virus, as described elsewhere (Sun et al., 2014). In contrast, Rajasethupathy et al. (2015) used a non-pseudotyped glycoprotein-deleted rabies virus which acts as a first-order tracer from the injection site (Wickersham et al., 2007). To test whether this strategy could provide off-target retrograde labelling, we made stereotaxic injections of a retrograde AAV vector (Tervo et al., 2016) into dorsal CA1 (figure 4A). Using this approach, in addition to dense neuronal labelling of CA1 pyramidal cells (Figure 4B), we saw labelling in the entorhinal cortex (figure 4F and 4G), as would be expected. However, similar to the extended figure 1 of (Rajasethupathy et al., 2015), we saw labelling of neurons directly above the injection site, and throughout both superficial and deep cortical layers in much of the brain (figure 4B; see also Extended Data figure 1a of Rajasethupathy et al., 2015). Additionally, we observed dense labelling of neurons in the anterior thalamic nuclei (figure 4E), which do not project to hippocampus proper (Mathiasen et al., 2020) and, importantly, we saw bilateral labelling in multiple subdivisions of prefrontal cortex, including the anterior cingulate cortex; it was most dense in the ipsilateral hemisphere (figure 4C). Without the selectivity of the pseudotyped rabies tracing approach from specific neuronal populations, we believe that the off-target retrograde labelling in this region was due to uptake of virus in axons passing through the cingulum and / or corpus callosum as well as along the path of the injection needle used.

**Figure 4:**
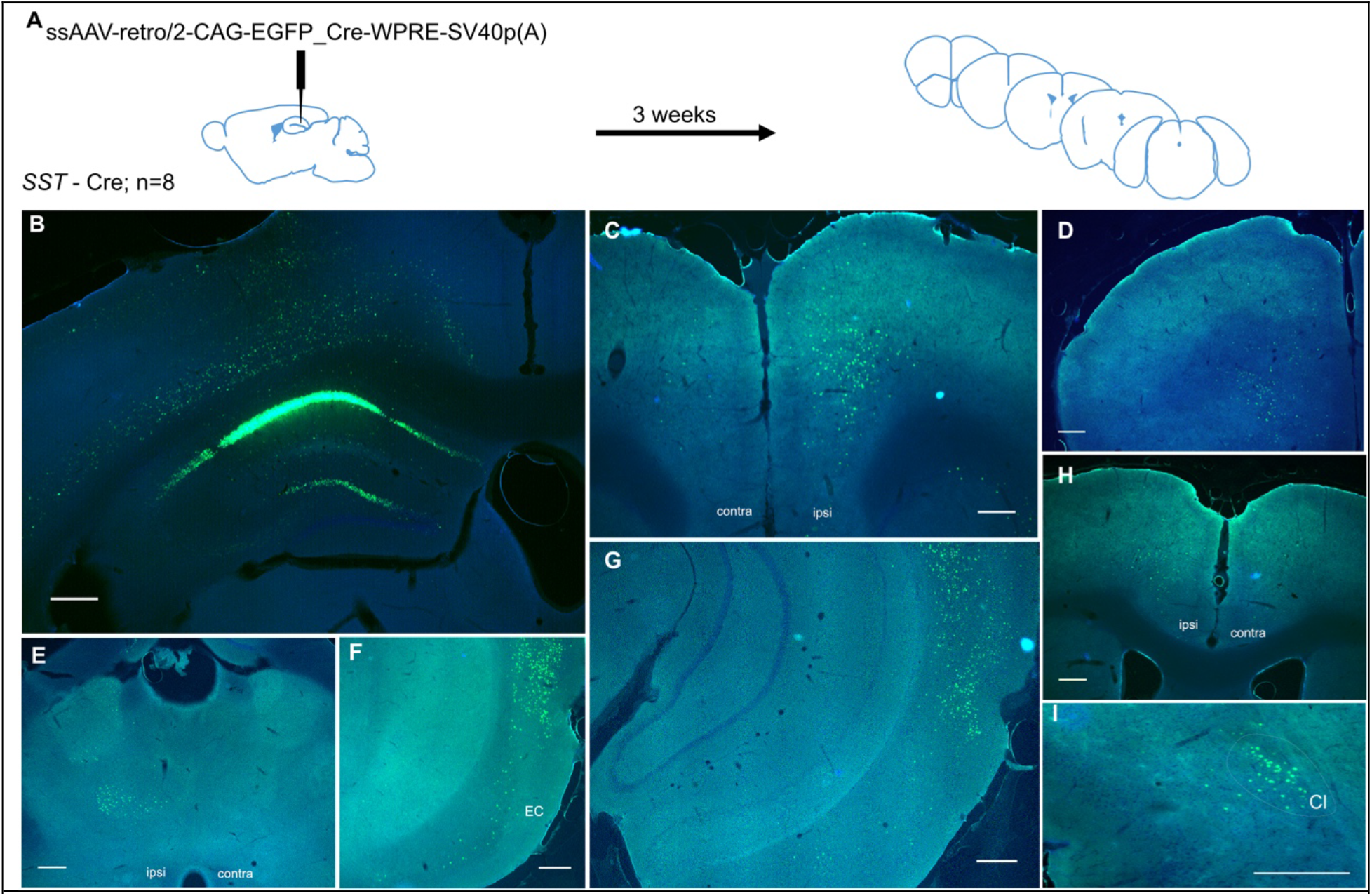
Retrograde tracing from dCA1 shows non-specific expression in multiple brain area, including both ipsi and contralateral anterior cingulate and prelimbic areas. **A)** outline of experimental method. Representative images of: **(B)** injection site dCA1, showing strong labelling in the hippocamus as well as some labelling in the overlaying cortical areas, **(C – D)** retrograde labelling in both ipsi- and contralateral hemispheres of prefrontal areas, **(E)** labelled cells present in ipsilateral thalamus, **(F – G)** entorhinal, perirhinal and ectorhinal cortices, **(H)** some labelled cells in ipsilateral and only a few in contralateral ACA, **(I)** retrograde labelled cells in ipsilateral claustrum. Scale bar 0.25 mm.

## Discussion

Here, we attempted to replicate the report of a novel projection from anterior cingulate cortex to hippocampus (Rajasethupathy et al., 2015) but failed to find any evidence of this projection in the mouse after examining a combination of tracing methods. We have also presented some technical issues that may have led to a misleading interpretation of the data in the original publication. We believe that it is important to report this failure to replicate, given that the existence of a direct anterior cingulate cortex projection to the hippocampus was published in a highly influential journal and, at the time of writing, the article has received hundreds of citations. Of those citations, we only found two other studies that reported looking for the anterior cingulate cortex to CA1 projection as part of a wider study (Fillinger et al., 2018; Wang and Ikemoto, 2016); both publications briefly noted that they failed to find evidence of this projection.

One recent publication (Bian et al., 2019) did present evidence of a monosynaptic projection from anterior cingulate cortex to CA1 (both dorsal and ventral) using retrograde AAV tracing as part of a study into prefrontal-hippocampal circuitry in contextual fear memory. Interestingly, this study found a far more sparse projection than that reported earlier (Rajasethupathy et al., 2015), with figure 5 of the Bian study revealing comparable ‘retrograde’ labelling in both M2 and cingulate areas to our results showing off-target labelling with the rAAV-retro vector. In that same study (Bian et al., 2019) the viral vectors failed to evidence of a connection from vHPC to anterior cingulate areas, despite such inputs having been described with more traditional axonal tracers (e.g. Hoover and Vertes, 2007; Jay and Witter, 1991). Such discrepancies highlight the potential idiosyncrasies of different tracer techniques.

### Stereotaxic coordinates and the borders of anterior cingulate cortex

Prefrontal cortex has been extensively studied in multiple model organisms, from mice and rats to humans and nonhuman primates. However, despite a large amount of homology across species, the shape of the prefrontal areas in the brains of different species is quite varied. These variations in shape, together with distinctions in the cytoarchitecture of the subdivisions of the prefrontal cortex have led to a lack of agreed nomenclature and clear boundaries of specific prefrontal regions (see Laubach et al., 2018 for a comprehensive analysis of discrepancies between authors). In this paper, we use the nomenclature and delineation from Vogt & Paxinos (Vogt and Paxinos, 2014) and the human homologies (Brodmann’s areas). A very helpful online unified anatomical atlas, combining the Franklin-Paxinos and the common coordinate framework from the Allen Institute of Brain Science, was also used cross check the anatomical borders of the prefrontal areas (Chon et al., 2019).

The injection spread of the anterograde tracing experiment, shown on figure 1b of the earlier study (Rajasethupathy et al., 2015) cleary includes the cingulate cortex, but also the M2 part of motor cortex is transduced by the virus; it is also possible that the transitional midcingulate area 24’ (Vogt and Paxinos, 2014) has been involved. Indeed, when using the reported coordinates, we again found significant spread of the virus in M2. Furthermore, looking at the projection pattern of the anterograde fibres in Rajasethupathy et al. (2015), it can be noticed that both hippocampi show fluorescent fibres, with the fibres in the contralateral hippocampus (right; figure 1b in their study) being brighter than in the ipsilateral hippocampus; the pattern, opposite to that in the case of the thalamus. Having used 250 nl of virus, as compared with 500 nl from Rajasethupathy et al. (2015) we still show the spillover neighbouring brain areas with these coordinates. Whilst the spillover of the virus itself cannot account for the apparent presence of axons from the cingulate cortex to CA1, nor the optogenetic activation of these afferents in that study, one potential explanation of a false positive could be that the relatively large volume of virus with a high titre injected close to the midline led to the leakage of virus beyond the intended injection site, resulting virus being taken up via axons traversing the cingulum bundle.

#### Concluding remarks

Here we report that we have been unable to find evidence of a monosynaptic projection from anterior cingulate cortex to hippocampus proper in the mouse, replicating findings reported over decades of rodent neuroanatomical literature but in stark contrast to two recent reports (Bian et al., 2019; Rajasethupathy et al., 2015). While we have presented some tentative explanations that could account for these discrepancies, we remain unable to provide a full account of why we and others failed to find evidence of this projection, should it exist. Our intention in reporting our findings is not to criticise the research carried out by colleagues, but to initiate a wider conversation about how we use genetic tools in systems neuroscience, and interpret the data. The hippocampal – prefrontal system is perhaps one of the most widely studied long-range circuits in cognitive neuroscience; that controversies still exist on the precise cellular connections between these two areas serves to highlight both the excitement and challenges that arise as modern genetic tools provide us with opportunities to study neural circuitry in detail that was unimaginable just 20 years ago.

## Acknowledgements

This work was supported by Biotechnology and Biological Sciences Research Council grant BB/P001475/1 (MTC). SY was a PhD student supported by Wellcome Trust grant 108891/B/15/Z. GMS and ESB were PhD students supported by grant MR/N0137941/1 for the GW4 BIOMED MRC DTP, awarded to the Universities of Bath, Bristol, Cardiff and Exeter from the Medical Research Council (MRC)/UKRI. Floxed TVA mice were kindly provided by Prof Dieter Saur at Technischen Universität München. We thank Dr E. Bubb and Dr A. Nelson (University of Cardiff) for access to relevant tracer data.

