## Supplementary data for "No evidence from complementary data sources of a direct projection from the mouse anterior cingulate cortex to the hippocampal formation"

### Supplementary material

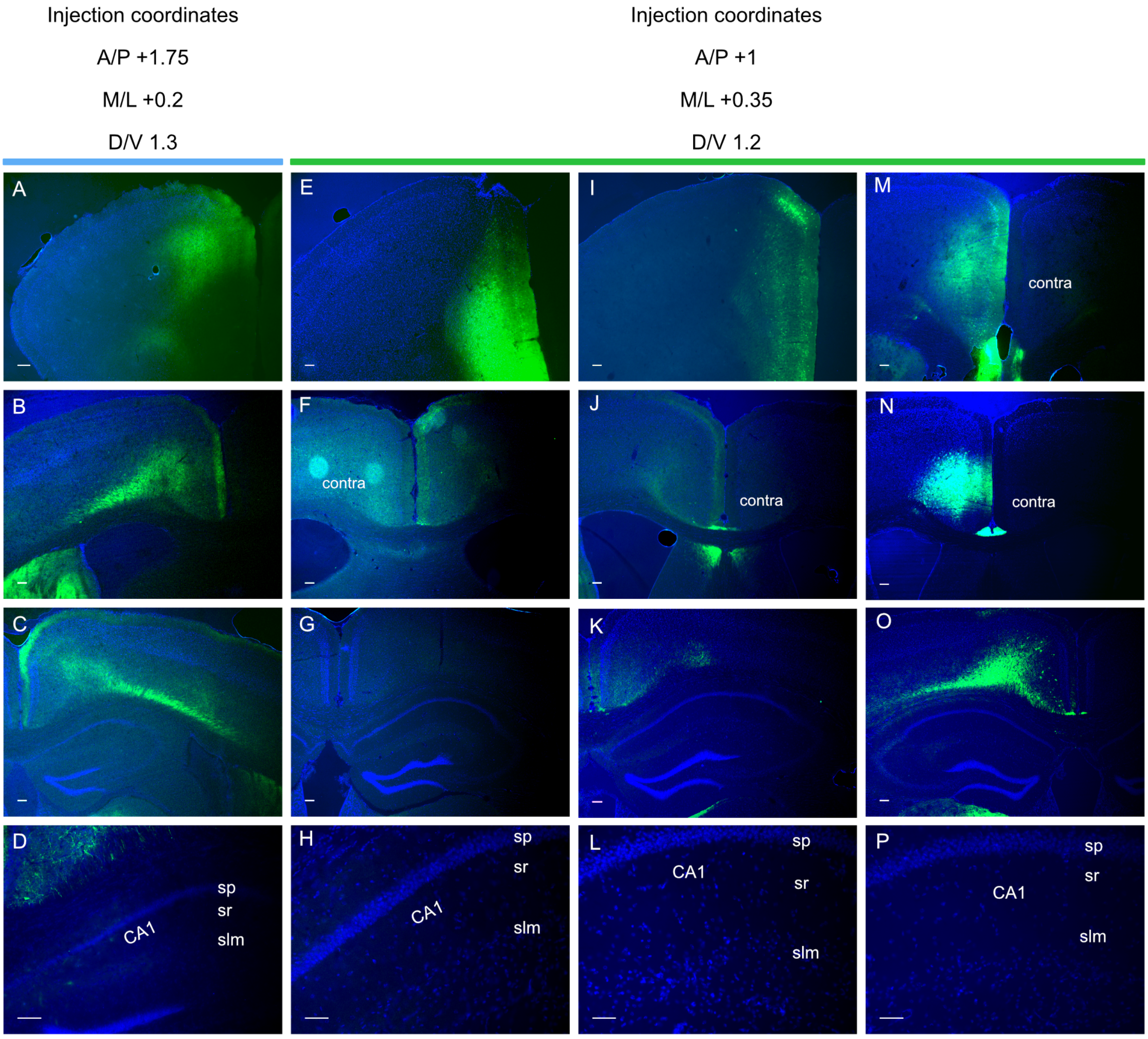


**Supplementary figure 1: Comparing viral spread in different injection coordinates of anterior cingulate area**. **A – D**, representative images of injection site with coordinates reported previously (A), spread into posterior part of ACA and other cortical areas (**B**), fluorescent signal present in many structures, but not in dorsal hippocampus (**C**), representative photo of dCA1 with no fluorescent signal present (**D**). **E – P**, representative images of 3 injections using in-house optimised coordinates to target anterior cingulate area. Similarly, fluorescent signal is present in other cortical areas (**F, J, N**) and absent in dorsal hippocampal sections (**G, K, O**), closer images of dCA1 with no fibres present (**H, L, P**).

**Supplementary table 1: Summary of all anterograde tracing AAV injections carried out.**

| | Coordinates (mm, relative to Bregma) | | | | | | | |
|---|---|---|---|---|---|---|---|---|
| Mouse ID | A/P | M/L | D/V | Injection site | Deep cortical projections (MCC and RSC) | Claustrum | Hippocampus | Thalamus |
| 1 | 1 | 0.35 | 1.2 | ACA/M2 | + | + | - | + |
| 2 | 1 | 0.35 | 1.2 | ACA/M2 | + | - | - | + |
| 3 | 1 | 0.35 | 1.2 | ACA/M2 | + | + | - | + |
| 4 | 1 | 0.2 | 1.3 | ACA | + | + | - | + |
| 5 | 1 | 0.2 | 1.3 | ACA | + | + | - | + |
| 6 | 1 | 0.2 | 1.3 | ACA | + | + | - | + |
| 7 | 1.75 | 0.20 | 1.50 | ACA | + | + | - | + |
| 8 | 1.75 | 0.20 | 1.50 | ACA | + | + | - | + |
| 9 | 1.75 | 0.20 | 1.50 | ACA | + | + | - | + |
| 10 | 1.75 | 0.20 | 1.50 | ACA | + | + | - | + |
| 11 | 1.75 | 0.20 | 1.50 | ACA | + | + | - | + |
| 12 | 1.75 | 0.20 | 1.50 | ACA | + | + | - | + |
| 13 | 1.75 | 0.20 | 1.50 | ACA | + | + | - | + |
| 14 | 1.75 | 0.20 | 1.50 | ACA | + | + | - | + |
| 15 | 1.75 | 0.20 | 1.50 | ACA | + | + | - | + |
| 16 | 1.75 | 0.20 | 1.50 | ACA | + | + | - | + |
| 17 | 1.75 | 0.20 | 1.50 | ACA | + | + | - | + |
| 18 | 1.75 | 0.20 | 1.50 | ACA | + | + | - | + |
| 19 | 1.75 | 0.20 | 1.50 | ACA | + | + | - | + |
| 20 | 1.75 | 0.20 | 1.50 | ACA | + | + | - | + |
| 21 | 1.75 | 0.25 | 1.75 | ACA | + | + | - | + |
| 22 | 1.75 | 0.25 | 1.75 | ACA | + | + | - | + |
| 23 | 1.75 | 0.25 | 1.75 | ACA | + | + | - | + |
| 24 | 1.75 | 0.25 | 1.75 | ACA | + | + | - | + |
| 25 | 1.75 | 0.25 | 2 | PFC/ACA | + | + | - | + |
| 26 | 1.75 | 0.25 | 2 | PFC/ACA | + | + | - | + |
| 27 | 1.75 | 0.25 | 2 | PFC/ACA | + | + | - | + |
| 28 | 1.75 | 0.25 | 1.75 | ACA | + | + | - | + |
| 29 | 1.75 | 0.25 | 1.75 | ACA | + | + | - | + |
| 30 | 1.75 | 0.25 | 1.75 | ACA | + | + | - | + |
